# Optimization of synthetic oscillatory biological networks through Reinforcement Learning

**DOI:** 10.1101/2023.11.19.567717

**Authors:** Leonardo Giannantoni, Alessandro Savino, Stefano Di Carlo

## Abstract

In the expanding realm of computational biology, Reinforcement Learning (RL) emerges as a novel and promising approach, especially for designing and optimizing complex synthetic biological circuits. This study explores the application of RL in controlling Hopf bifurcations within ODE-based systems, particularly under the influence of molecular noise. Through two case studies, we demonstrate RL’s capabilities in navigating biological systems’ inherent non-linearity and high dimensionality. Our findings reveal that RL effectively identifies the onset of Hopf bifurcations and preserves biological plausibility within the optimized networks. However, challenges were encountered in achieving persistent oscillations and matching traditional algorithms’ computational speed. Despite these limitations, the study highlights RL’s significant potential as an instrumental tool in computational biology, offering a novel perspective for exploring and optimizing oscillatory dynamics within complex biological systems. Our research establishes RL as a promising strategy for manipulating and designing intricate behaviors in biological networks.

## I. Introduction

Synthetic biology represents the convergence of biology and engineering, aiming to design and construct novel biological entities or reconfigure existing ones for specific purposes [1]. A key application in this field is biological computing. Unlike traditional silicon-based computing systems, biological ones use cells, proteins, and DNA to engineer biological motifs for computational tasks [2]. With their small size and energy footprint, these bio-based systems present unique advantages over silicon-based computation [3]. They also hold the potential for self-replication and self-assembly [4]. However, designing large, intricate, and scalable synthetic biological systems presents considerable challenges [5], especially in maintaining a consistent temporal framework [6]. In this context, synthetic oscillators Figure 1, bioengineered to produce periodic signals through molecular concentration fluctuations [7], are vital. They mirror natural rhythmic processes using genes and, just as quartz crystal clocks underpin time operations in silicon-based computers, they are essential for timing [7]–[9] and information representation [10] in biological computing. Despite significant progress in understanding these systems, linking the complex behaviors of naturally occurring oscillators to their efficient and robust analogous in synthetic biology remains a challenging area of investigation [11], [12].

**Fig. 1.**
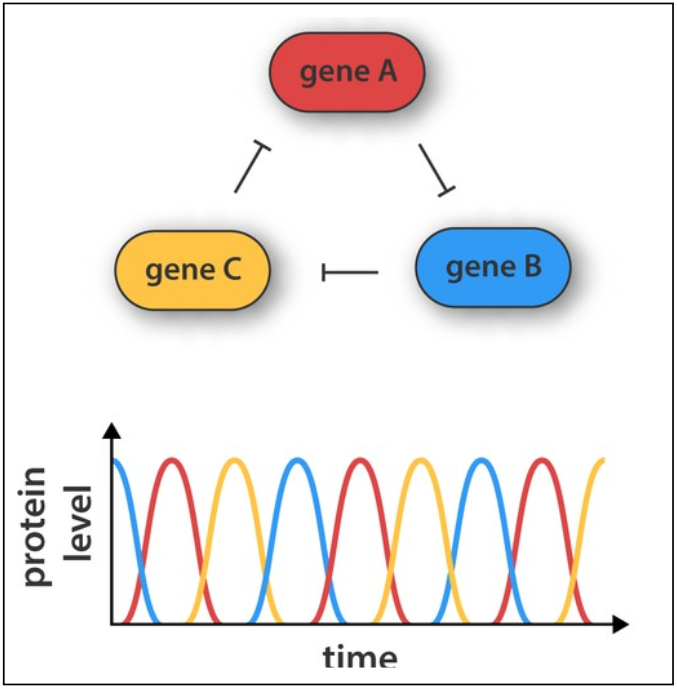
Synthetic gene network containing three genes and generating oscillations. Each gene is translated into a protein product, repressing the activity of another gene in the network (as indicated by the arrows) [27].

Natural rhythms have been studied for a long time, and much has been understood about the behavior of simple units of biological computation. Historically, feedback loops have been recognized as central to the dynamics of biological systems [13], [14]. Both negative and positive feedback loops play crucial roles in oscillations and adaptability [15]– [While negative feedback loops are vital for oscillations [15], [18], robust positive feedback loops are essential to period adaptability [19]. A straightforward way to produce persistent oscillations has also been demonstrated by employing a negative feedback loop with a time delay [20], [21].

Although oscillations can occur even in systems comprising only a tiny number of proteins [22], oscillatory dynamics in nature usually arise from the intricate interplay of many feedback loops [23] that produce complex qualities such as transients, history-dependence and after-effects, frequency scaling, temperature dependency [24].

The diversity and complexity of natural oscillators, from calcium oscillations in cells to neural oscillations at macroscopic scales, underscore the rich dynamical behavior of these systems [11], [25]. However, understanding and harnessing these oscillations in synthetic networks is challenging due to their non-linearity, high dimensionality, and potential discontinuities [23].

Traditional methods for the design and analysis of such networks can be computationally intensive, as the inherent characteristics of larger networks do not offer easy mathematical characterization [12] nor a smooth landscape, generating a vast design space of possible structures and parameter values [11], [26].

This work introduces an Optimization via Simulation (OvS) approach using Reinforcement Learning (RL) [28] (RL-OvS) for designing synthetic biological oscillators, addressing the challenges posed by molecular noise and system complexity. RL is particularly suited for this task because it navigates complex, high-dimensional spaces without explicit gradient information. This significantly differs from traditional computational methods, which often rely on predefined rules and strategies. RL not only adapts and learns from the system’s feedback but also builds a mathematical model representing the relationship between design parameters and system behavior. This capability of learning and adapting in real-time to dynamic inputs, such as molecular noise, positions RL as a novel and efficient tool in the computational biology domain, particularly for exploring and optimizing oscillatory dynamics. Our approach aims to demonstrate the potential of RL in this field, offering a new perspective compared to traditional algorithms and underscoring its relevance and applicability in synthetic biology and biocomputing.

## II. Background

The pursuit of automated design and optimization in synthetic biology represents a frontier in integrating engineering principles with biological complexity. Traditional efforts have primarily focused on leveraging empirically established design principles to confirm experimental viability and propose potential structures. However, such approaches often limit the diversity and number of constituent devices they can effectively manage. Furthermore, a critical gap in these studies is the lack of predictive modeling capable of facilitating more efficient Design Space Exploration (DSE) and uncovering novel design principles and motifs.

In genetic oscillatory circuit design, a spectrum of techniques has been employed. These range from exhaustive searches and optimization-based techniques to more sophisticated control theory and rule-based methods. These advanced methods necessitate a deep understanding of circuit properties and their relationship to network topology [29].

For instance, one approach in this area has involved integrating rule-based design with Monte Carlo simulated annealing algorithms to optimal topologies from a library of standard biological components [30]. This approach, while versatile, relies heavily on a predetermined set of rules and is limited by the static nature of these rules. Similarly, evolutionary algorithms have been explored to create circuits exhibiting rhythmic or switching behaviors [31], [32]. These methods, although effective, tend to face scalability issues and become computationally intensive as the complexity of the circuits increases. The Mixed-Integer Nonlinear Programming (MINLP) framework, as proposed in various studies [33]–[36], also suffers from high computational demands, particularly when dealing with complex circuit designs.

Evolutionary algorithms have also been used, as seen in [31], which focuses on identifying optimal circuits exhibiting specific rhythmic or switching behaviors, and in [32], where evolutionary algorithms are used in conjunction with multiobjective optimization. Despite their effectiveness, these algorithms face scalability challenges, mainly when applied to more complex circuit designs. The MINLP framework, as proposed in various studies [33]–[36], also suffers from high computational demands, mainly when dealing with complex circuit designs. In particular, the choice in [34]–[36] of synthesizing biological oscillators among those combinatorially obtained using three devices from a library of 44 [37], severely limits from the start the complexity of the designs.

In [38], an attempt is made to integrate classical optimization methods with Machine Learning (ML) to accelerate the discovery of gene circuits. This approach, which models networks as systems of coupled Ordinary Differential Equations (ODEs), proved more efficient than evolutionary algorithms and scalable for networks with up to nine nodes [29]. Nonetheless, this methodology, like the others, does not inherently capture the causal relationships within the network.

Bifurcation analysis, a technique employed to optimize parameters for inducing Hopf bifurcations or turning point bifurcations, offers insights into the dynamic behavior of oscillatory and bi-stable circuits [39]. In [40], this approach is used with an evolutionary algorithm that progressively tunes randomly generated network structures. While both methods can be rapidly executed, they still do not facilitate learning causal relations.

This overview of the existing methods underscores the need for approaches that can not only navigate the complex parameter space of synthetic oscillatory systems but also learn and adapt based on system behavior. RL stands out as a promising candidate due to its ability to interact with and learn from the system iteratively, a feature critically missing in most traditional methods. The potential of RL to understand and leverage the causal relationships in biological networks presents a novel approach to the challenges of designing robust and efficient synthetic oscillators.

## II. Methods

### A. Materials

This study focuses on selecting and optimizing biological networks structured as oscillators. The primary objective is to adjust the system parameters to achieve sustained oscillations despite molecular noise. These parameters encompass the initial concentrations of both floating and boundary species and their rate constants. To realistically simulate molecular noise, we initialize these parameters by drawing from a uniform distribution that spans a wide range of values that reflect biological variability.

Central to our approach is optimizing floating species, targeted for their oscillatory potential, as opposed to boundary species that typically act as molecular sources or sinks. This distinction is vital in our strategy to harness and control oscillatory dynamics within these networks.

The biological networks we optimize are sourced from existing literature, where they are represented in two primary forms: as systems of stoichiometric equations or through Systems Biology Markup Language (SBML) notation [41]. These representations are then converted into ODE systems, forming the basis for our optimization process. The optimization is conducted via simulation, where the RL algorithm iteratively adjusts parameters to improve the system’s oscillatory behavior. This process is rigorously evaluated using principles from bifurcation analysis, focusing on identifying and reinforcing the conditions that lead to sustained and stable oscillations.

Further details on these tools and concepts and the specific methodologies employed in our optimization process will be elaborated in the following sections. The efficacy and outcomes of our approach are demonstrated through two distinct case studies, which are discussed in section IV.

### B. Modeling and analysis

This section outlines the formalism used to represent and analyze models of oscillators, which forms the basis for our evaluation and scoring algorithm as described in subsection III-C-Reward.

Biochemical networks are modeled using systems of stoichiometric equations or ODEs. These mathematical representations are crucial for capturing the dynamic interplay of multiple biological processes and states. ODEs, in particular, provide a powerful tool for describing how the concentrations of different species in a biological system evolve.

Stoichiometry refers to the relationship between the reactants and products in a reaction, presented in a balanced equation that reflects the conservation of mass. For instance, consider a simple biological reaction represented by the following stoichiometric equations:

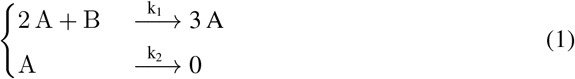

In this system (Equation 1), the first equation describes a reaction converting species *B* into *A* with a rate constant *k*_1_, while the second represents the degradation of *A* with a rate constant *k*_2_. This can be equivalently described by the following ODE system:

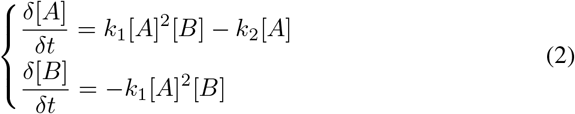

Equation 2 describes the rate of change in the concentrations of compounds [*A*] and [*B*] over time, where [*A*] and [*B*] represent the concentration of species *A* and *B* in the system, respectively. The coefficients *k*_1_ and *k*_2_ dictate how fast these reactions occur. In the first equation, the first term implies that the reaction rate depends on the square of [*A*] and linearly on [*B*], while the second term is directly proportional to [*A*]. Similarly, for the second equation.

The core of our analysis lies in understanding how these systems exhibit oscillatory behavior, often characterized by a Hopf bifurcation. A Hopf bifurcation marks a critical transition from a steady state to a periodic oscillating state, depending on the values of specific parameters in the system. This is determined by analyzing the eigenvalues of the Jacobian matrix of the ODE system, which provides insights into the local stability of the system around a fixed point. For example, if the real parts of a complex pair of eigenvalues of the Jacobian cross the imaginary axis, a Hopf bifurcation occurs, indicating the onset of limit cycle behavior or sustained oscillations in the system. Furthermore, the eigenvalues’ imaginary parts correspond to the oscillations’ frequency. The imaginary axis’ crossing marks where these inherent oscillations manifest in the system’s dynamics [42], [43].

Consider a biological system described by *N* nonlinear, coupled, ordinary differential equations, whose state is described by *N* variables *X*_*i*_, *i* = 1 : *N*, and by the parameters

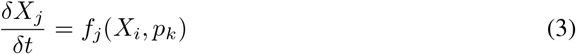

where *f*_*j*_(*X*_*i*_, *p*_*k*_) are, in general, nonlinear functions of all the *X*_*i*_, *p*_*k*_. The eigenvalues of the Jacobian (*∂f*_*i*_*/∂X*_*j*_) determine the system’s stability at a steady state. If 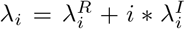 are the complex eigenvalues of the Jacobian for a particular steady state, then instability will result if any of 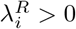. If a complex pair of eigenvalues becomes purely imaginary, then we get a Hopf bifurcation, and the system will exhibit limit cycle behavior [39].

In our RL algorithm, this understanding of Hopf bifurcations plays a pivotal role. The reward computation described in subsection III-C-Reward is directly influenced by the system’s proximity to a Hopf bifurcation, as it signifies the achievement of desired oscillatory dynamics. Thus, the RL algorithm is designed to manipulate the system parameters in a way that drives the system towards or maintains it in this critical oscillatory state. This approach allows the algorithm to learn strategies that effectively tune the biological networks towards stable, sustained oscillations, a key objective in our study of synthetic biological oscillators.

### C. Optimization via simulation

Our proposed Reinforcement Learning Optimization via Simulation (RL-OvS) framework, as depicted in Figure 2, is designed to automate exploring the design space for synthetic biological oscillators.

**Fig. 2.**
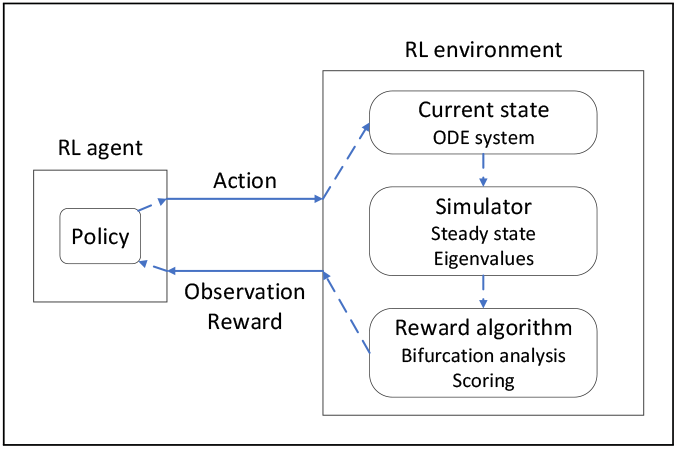
RL-OvS framework. The agent selects actions based on observations and rewards, while the environment provides feedback and updates the state.

This framework (Figure 2) comprises two interacting components, an *agent* and an *environment*, each playing a crucial role in the optimization process.

The *agent* is tasked with exploring the design space and identifying optimal configurations of biological networks, i.e., the initial concentration of floating and boundary species in the ODE system, by observing the systems’ behavior and acting upon it, mediated by the *environment*. It operates by receiving and interpreting data about the current state of the system, known as *observation*, and a *reward* that quantifies the system’s performance. Based on these inputs and its evolving *policy*, the agent makes decisions, or *actions*, aimed at optimizing the system’s oscillatory behavior.

Conversely, the *environment* is responsible for implementing the actions taken by the agent. It modifies the underlying ODE system according to the agent’s directives, simulates the resulting dynamics, and evaluates the outcome. The evaluation is based on bifurcation analysis, specifically focusing on the system’s proximity to a Hopf bifurcation, a critical juncture where the system transitions between different dynamic behaviors. The outcome of this evaluation is fed back to the agent as a *reward*, along with updated *observation* data, informing the agent’s subsequent decisions.

These enable the agent to update its *policy* and choose the following *action*. The iteration of these steps lets the agent learn an optimal *policy*, a strategy for solving the problem.

This interactive process is structured into discrete steps, each constituting a part of an *episode*. Within each episode, the agent iteratively refines its policy, balancing the *exploration* of new actions with the *exploitation* of known beneficial strategies. This process is crucial for the agent to learn and adapt its approach to maximize the cumulative reward and discover an optimal policy. The episodic nature of this learning process also ensures that the agent gains experience from a wide range of scenarios, contributing to the robustness and generalization of the learned policy. Indeed, multiple episodes are essential in RL to let agents try different strategies. During early episodes, the agent might make suboptimal decisions that lead to undesirable outcomes. By experiencing the consequences of its actions over multiple episodes, the agent can adjust its policy to make better decisions in the future. Multiple episodes are also needed to ensure robustness and generalization through various experiences. As with biological systems, this aspect is essential in dynamic and potentially noisy environments. By averaging the learning across multiple episodes, the agent can mitigate these aspects and learn a more stable policy [28].

RL ‘s capability to explore and exploit the action space allows it to navigate the vast possible configurations of biological networks, balancing the discovery of novel control strategies with the refinement of known beneficial actions. This adaptive learning process aligns closely with biological systems’ inherent variability and adaptability.

The core of our RL-OvS framework is a highly interactive environment that challenges and guides the RL agent’s learning process. Implemented in Python, this environment utilizes Gymnasium [44] for a standardized RL interface, facilitating integration with various RL algorithms. Our environment specifically models the biological network as an ODE system, simulating and evaluating its behavior to provide targeted feedback to the agent.

For simulating oscillatory models, we leverage Tellurium [45], a versatile tool that supports both SBML and Antimony formats. This allows us to handle a range of computational models, from simple stoichiometric systems to more complex dynamic networks, and to conduct both deterministic and stochastic simulations. This flexibility is crucial for accurately capturing biological oscillators’ diverse dynamics.

#### State and observation

Within our framework, the state of the environment encapsulates the current conditions of the modeled biological system. This includes the concentrations of various species and additional parameters indicative of the system’s dynamic state. The observation that the agent receives is a distilled version of this state, containing all essential information needed to make informed decisions. This could range from current species concentrations to system stability or instability indicators.

#### Action space

An action is an agent’s specific decision or move that affects the environment’s state, aiming to maximize its cumulative reward over time. The agent’s actions directly influence the biological system’s parameters, with options to increment, decrement, or multiply these parameters within a predefined range. This range is carefully chosen to ensure biological relevance while allowing sufficient parameter space exploration.

#### Agent’s Learning Process

Employing the Proximal Policy Optimization (PPO) algorithm [46] from the Stable Baselines3 library [47], our agent iteratively refines its policy based on the feedback received. PPO, a policy gradient method, operates by optimizing a surrogate objective function to ensure that the new policy does not divert too drastically from the old policy, thus preventing overly aggressive policy updates that can destabilize learning. PPO ‘s advantage thus lies in its ability to balance exploration with exploitation, ensuring stable and consistent learning even in the complex and often unpredictable landscape of biological systems.

#### Reward mechanism

The reward mechanism is central to the agent’s learning and is designed to guide the agent towards desirable system states. Based on a bifurcation analysis of the system’s eigenvalues, the reward computation serves two primary purposes. Firstly, it provides positive reinforcement for states approaching or achieving a Hopf bifurcation (Equation 4), indicative of desired oscillatory behavior. Secondly, it penalizes states that either deviate significantly from this behavior or lead to system instability (Equation 5). This dual mechanism ensures that the agent is consistently nudging the system towards stable and sustained oscillations, which are hallmarks of functional biological oscillators.

Given a system with *n* eigenvalues, Equation 4 assigns a positive score for each *n*_*th*_ eigenvalue indicating the presence of a putative Hopf bifurcation, that is when the real part 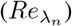 of the eigenvalues goes to zero. If at the same time that *n*_*th*_ eigenvalue has an imaginary part 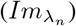, the assigned score is 1. The latter condition is to differentiate systems having no imaginary components, thus presenting a turning point or a static bifurcation point [39], [40], and awarding them a penalty (Equation 5). Otherwise, when 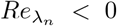 (subsection III-B), the assigned score is 0.1, weighted by the *gap* between the real and imaginary parts of the eigenvalue to take into account dominance. This smaller score is meant to encourage approaching the bifurcation. Ideally, a system with no damping would have purely imaginary eigenvalues 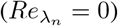.

Equation 5 computes penalties. It assigns a large negative score (−100) when a steady state (*ss*) solution does not exist, the Jacobian matrix is non-invertible, or the systems’ eigenvalues have no imaginary parts. Otherwise, it assigns 10 for each 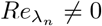.

Finally, the reward (Equation 6) is the sum of these two contributions, one encouraging the approach to an oscillating state, the other penalizing configurations that deviate from it, making the system unstable or too far from a Hopf bifurcation.

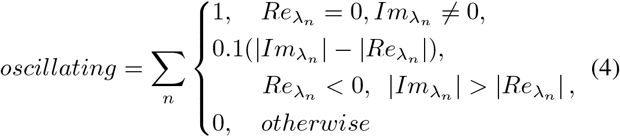

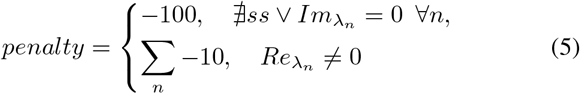

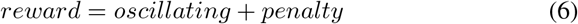

## IV. Results

Our investigation into the capabilities of the RL-OvS framework in optimizing biological oscillators was conducted through two distinct case studies. These case studies were chosen to highlight the versatility and potential of our approach in navigating the dynamics typical of biological oscillator models.

### A. Setup

The setup for our experiments was carefully designed to provide an effective testbed for the PPO learning algorithm (see section III). To this end, the following key parameters were carefully chosen [46], [47]:

- Minibatch size: number of episodes used in one iteration of the gradient descent update. Smaller values can lead to noisier gradient updates, while larger ones provide more stable updates but require more memory.
- Rollout buffer size: number of time steps for an agent to interact with the environment before the data is used for learning. The data collected is stored in a buffer. The buffer size can impact the quality and diversity of experiences for each update.
- Learning rate: step size at each iteration while moving towards a minimum of the loss function. A high learning rate can cause the algorithm to converge faster but might overshoot the optimal solution or oscillate. A low learning rate can make the convergence slower but might be more precise.
- Total timesteps: The total number of samples (i.e., environment steps) to train on.

As with most RL algorithms, tuning these parameters appropriately is crucial for PPO to balance computational efficiency with the quality of learning. We adopted a batch size of 32, a rollout buffer of 640 steps, a learning rate of 0.001, and 100 timesteps per episode, spanning 400 iterations.

Furthermore, we incorporated random initialization of model parameters at the beginning of each episode. This approach encouraged diverse exploration of the parameter space and mirrored the variability and unpredictability inherent in biological systems, enhancing the robustness and generalizability of the learning outcomes.

The computational resources and time required to run the RL algorithm for each case study varied notably, reflecting the distinct complexities of the biological models. The first case study, which involved the Edelstein relaxation oscillator (Edelstein relaxation oscillator), ran for approximately 6 hours on an HP 855 G8 laptop equipped with an AMD Ryzen 7 5700U CPU @ 1.8 GHz and 32 GB RAM. This duration was primarily due to a significant proportion of unfeasible solutions that were filtered out, such as those lacking a steady state. In contrast, the second case study, focusing on the Otero repressilator (Otero repressilator), ran for approximately 8 hours on the same laptop configuration. This difference in execution time between the two case studies highlights the diverse computational needs relative to the biological models’ complexity. It suggests a potential area for future refinement in simulation length and computational efficiency.

### B. Case study 1 - Edelstein Relaxation Oscillator

The first case study focused on the Edelstein relaxation oscillator model, as outlined by Seno et al. in 1978 [48]. This model describes biological systems that undergo rapid transitions between two states, with a slow evolution in one state and a rapid *relaxation* or transition to the other. This behavior can be likened to neuron firing, where an immediate discharge follows a slow buildup of electrical potential. Besides its relevance in biology, through this example, we are positioning our study to make direct comparisons with other research, such as [40].

We observed a clear progression in the learning process through a series of simulations (synthetically illustrated in Figure 3 A-F). Early stages of the learning process (panels A and B) demonstrated a lack of sustained oscillations, indicating initial challenges in system tuning. However, as the learning progressed, the algorithm gradually guided the system towards more regular oscillatory behavior (panels C, D, and E), culminating in panel F, where three out of four species exhibited oscillatory patterns, albeit with some damping. While indicative of our approach’s potential, this progression also highlights areas for further refinement, particularly in tuning hyperparameters and improving reward allocation strategies.

**Fig. 3.**
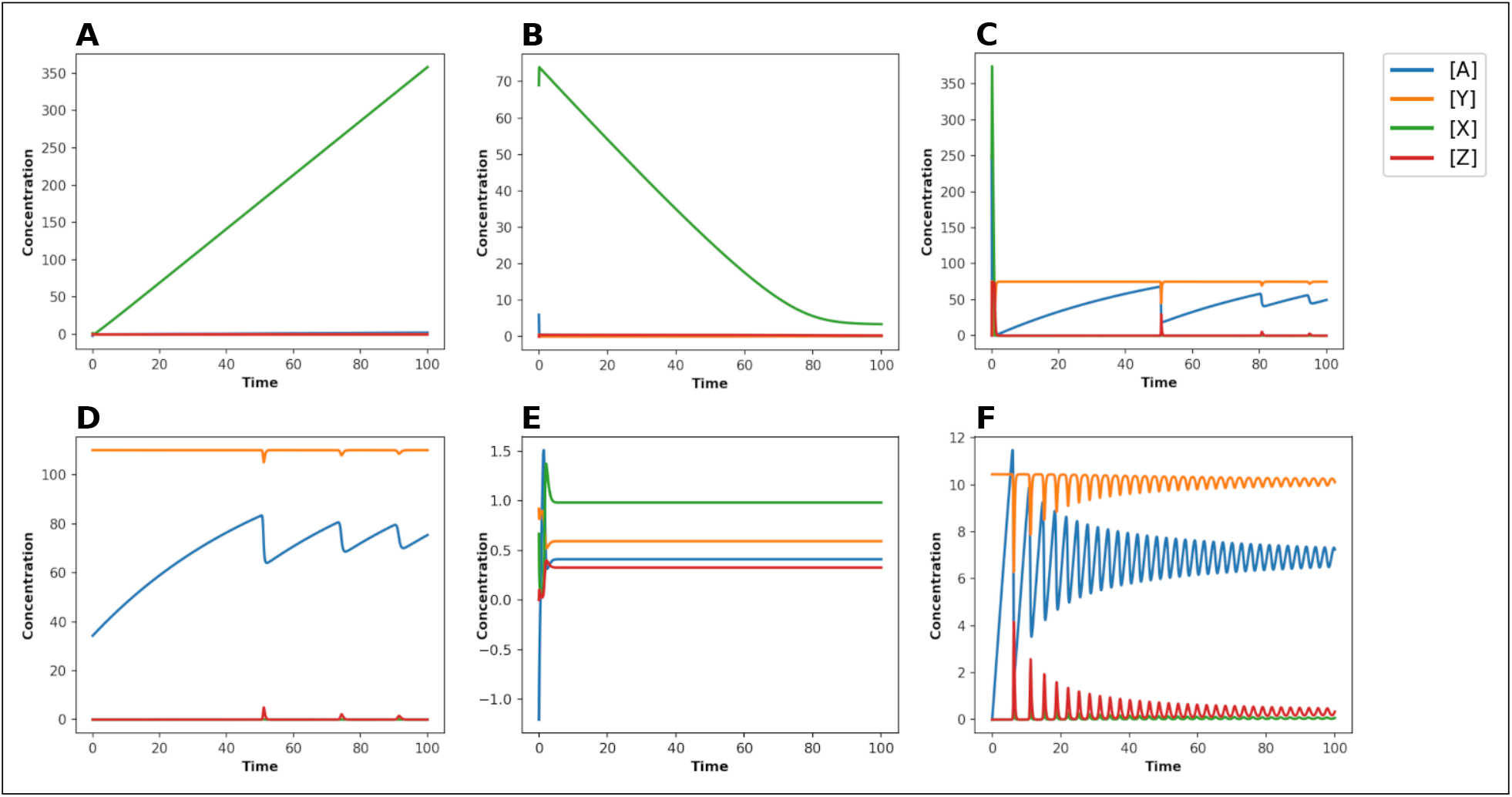
Edelstein relaxation oscillator. Evolution over time (from A to F) of the optimized circuits, showcasing the progression from non-oscillatory to oscillatory yet damped behavior (F).

### C. Case study 2 - Otero repressilator

The second case study involved the *Otero repressilator*, an oscillator model presented in [35]. Its structure, depicted in Figure 4, was found by the authors using a library of biological parts augmented with the degradation of the bound repressor and has a similar topology to the original repressilator system by Elowitz [7].

**Fig. 4.**
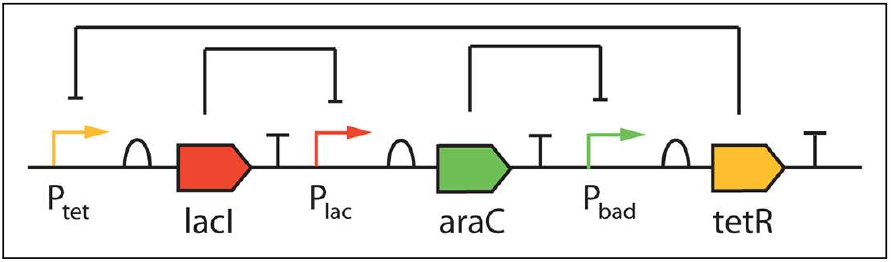
Case study 2 - Structure of the Otero repressilator [35].

As shown in Figure 5, the results were particularly promising and more encouraging than the Edelstein relaxation oscillator case study.

**Fig. 5.**
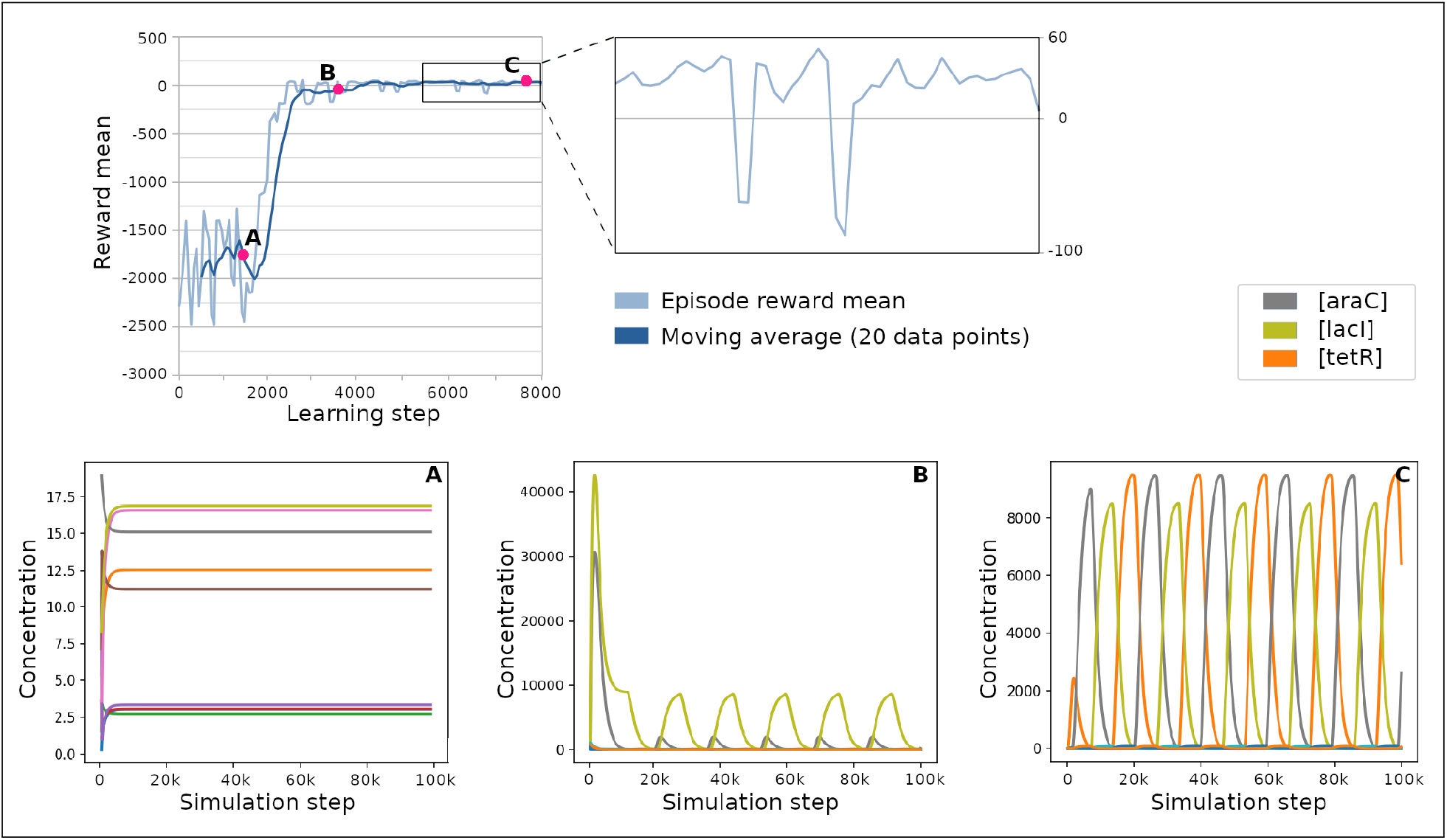
Edelstein relaxation oscillator. Episode reward average over the learning updates and three networks corresponding to different optimization stages. The average reward changes through exploration steps, hovering around -2000 for 2000 iterations, then reaches 0 and keeps growing slowly but steadily.

The upper-left panel in Figure 5 shows a line chart of the evolution of the average episode reward over 8000 learning algorithm update steps, with the episode reward mean computed at the same time as the update for each one of the *rollout_buffer_size/batch_size* = 640*/*32 = 20 mini-batches, times 400 iterations. The darker line is a moving average of 20 such updates. This panel shows that the mean reward hovers around a low value of approximately -2000 for an extended initial period.

However, a marked improvement in the reward pattern was observed after 2000 learning steps, indicating a significant shift in the system’s dynamics towards more favorable states. This favorable trajectory sustains throughout the remainder of the iterations, with a gentle yet persistent ascent observed. On the top right, a zoomed-in section of the last 2000 updates highlights the rewards’ consistency above zero, with only a few deviations.

The lower section in Figure 5 shows simulation results of the oscillator networks. On each chart (A-C), the horizontal axis represents simulation time, and the vertical axis represents species concentration. The species under optimization in this oscillator are the three proteins *araC, lacI*, and *tetR*. These three panels capture the oscillator’s state around 2000 (A), 4000 (B), and 8000 (C) learning updates. Their corresponding position is marked with red dots on the reward curve on the top-left chart.

Detailed analysis of the oscillator’s behavior at these stages reveals a progressive shift towards a synchronized oscillatory state among all involved species. As expected from the episode reward mean, the circuit simulated in panel A lacks oscillations. In contrast, the middle image (panel B) captured around 4000 iterations shows explicit oscillatory behaviors in two of the three species, i.e., *araC*, and *lacI* proteins. After an initial spike, these species exhibit sustained oscillations that do not appear to dampen over time. However, the third species – namely, *tetR* – remains non-oscillatory to casual observation. By the final stage, around 8000 learning updates, the third circuit (panel C) portrays all three species in synchronized oscillation without pronounced spikes and damping. This synchronization, particularly evident in the later stages, not only validated the capability of our algorithm to adapt and learn but also underscored its potential in steering complex biological systems toward specific dynamic states.

## V. Conclusions

This study investigated the potential of RL in optimizing synthetic biological networks’ dynamic behavior, specifically focusing on oscillatory dynamics and controlling Hopf bifurcations in ODE-based systems. Our findings reveal that RL demonstrates adaptability to biological systems modeled as ODEs, albeit with challenges in achieving persistent oscillations. While promising, the Edelstein relaxation oscillator case study indicated areas for refinement. Additionally, our RL-based approach for bifurcation analysis shows computational efficiency, although with longer execution times than traditional methods. This highlights opportunities for optimization and speed enhancements in future iterations, emphasizing RL’s potential in dynamic systems.

In conclusion, our study reinforces the notion that RL holds significant potential as a tool in computational biology, particularly for addressing the multifaceted challenges presented by complex biological systems. As we move forward, a clear pathway exists for applying RL in designing biological networks with more precise dynamic behaviors. Future research will focus on refining the reward computation mechanisms and exploring the use of multiple RL agents working collaboratively within an environment. This collaborative approach could facilitate the assembly of more complex systems from basic biological motifs, pushing the boundaries of what can be achieved in synthetic biology.

## VI. Data and code availability

The source code and data to reproduce the results presented in this paper and to apply the same procedure to other datasets are available from the Open Access repository https://github.com/smilies-polito/RLoscillators.

## Notes

https://github.com/smilies-polito/RLoscillators

### Competing Interest Statement

The authors have declared no competing interest.

